# Ultrastructural comparison of human kidney organoids and human fetal kidneys reveals features of hyperglycemic culture

**DOI:** 10.1101/2023.03.27.534124

**Authors:** Anika Schumacher, Virginie Joris, Martijn van Griensven, Vanessa LaPointe

**Affiliations:** Department of Cell Biology–Inspired Tissue Engineering, MERLN Institute for Technology-Inspired Regenerative Medicine, Maastricht University, Maastricht, The Netherlands

**Keywords:** Nephrogenesis, kidney organoid, diabetic nephropathy, transmission electron microscopy, GSK3β, glycogen

## Abstract

Induced pluripotent stem cell (iPSC)–derived kidney organoids have the potential for a large variety of applications. However, they do not persist long in culture, for which reasons are still unclear. Furthermore, their morphological maturation, an essential feature for kidney function, has not been sufficiently assessed. Kidney organoids are transcriptionally much alike end-of-first-trimester fetal kidneys and present many of the same cell types. From large transmission electron microscopy tilescans of specific regions of interest, we compared the ultrastructures of iPSC-derived kidney organoids at various timepoints to human fetal kidneys of the first trimester. Unlike healthy fetal kidneys, large glycogen deposits developed over time in all organoid cell types, but particularly in podocytes and in chondrocytes, one of the off-target populations that contaminate the culture. Deeper investigation showed that glycogen synthase kinase 3b (GSK3β) levels and activation were diminished over time, correlated with the accumulation of glycogen. Activated YAP was strongly expressed and large lipid droplets accumulated over time in proximal tubules. Accordingly, EGFR signaling increased significantly over time. Mitochondria in glomeruli and tubules contained few or no cristae, indicating mitochondrial damage. Together these features are known for hyperglycemic cultures and diabetic nephropathy. Measuring the glucose concentration in the kidney organoid culture medium showed a concentration of 2.94 g/mL, which is considered an elevated, pre-diabetic–like concentration *in vitro*. In summary, our ultrastructural assessment of iPSC-derived kidney organoids using an age-matched fetal human reference allowed the evaluation of cellular morphology, and we identified intracellular features that can inform the cellular state, which is particularly important while physiological testing of organoids is limited.

**Translational Statement:** Kidney organoids hold promise as a future treatment for patients with end-stage kidney disease. The engineering of kidney organoids with correct and healthy morphology *in vitro* is therefore essential, to guarantee functionality after transplantation. The present study provided deeper insights into the structural organization and ultrastructure of cells in kidney organoids compared to age-matched human fetal kidneys. Accordingly, we found several features in the regular kidney organoid culture, which are known to occur in hyperglycemic cultures and diabetic nephropathy, indicating that the current medium composition may be inducing pathological cellular phenotypes. This study therefore creates a better understanding of current limitations in the kidney organoid culture, increases knowledge of their function and cellular organization, and sets the foundation for further research to create advanced organoids.

## Introduction

In recent years, self-organizing microtissues like kidney organoids have shown great potential for disease modeling, drug testing and future transplantation to restore the function of diseased or damaged kidneys. To assure functionality upon transplantation, kidney organoids must faithfully replicate key features of the human kidney. Depending on the application, correct cellular identity, organization, morphology and functionality are essential. However, to date, human pluripotent stem cell–derived kidney organoids stagnate in growth after approximately three weeks (D7+25) and transcriptionally mature no further than fetal kidneys at the end of the first trimester of gestation.(1)

Current kidney organoid literature focuses on assessing protein and gene expression to describe the phenotype development. With this, the presence of approximately 5–6 renal cell types, excluding a number of mesenchymal/stromal populations, has been confirmed.(2–4) Compared to fetal kidneys of similar age, the cellular diversity is incomplete.(5) Early evidence suggests that these organoids are also morphologically immature, even after transplantation.(6) However, research on the morphology of kidney organoids in general is scarce and it is unclear how well kidney organoids morphologically replicate fetal kidneys of similar transcriptional age.

Ideally, kidney organoid ultrastructure would be assessed in comparison to human fetal kidneys. In our recent work, we described the (ultra-)structural development of human embryonic and fetal kidneys at Carnegie stage 20 until post-conceptional week (PCW) 12, a similar timepoint to the endpoint of the kidney organoid culture.(7) These findings enable the assessment of the structural maturity of kidney organoids and pinpoint divergent features that would need improvement to generate organoids more similar to human (fetal) kidneys.

There are numerous morphological features that are essential to be correctly replicated to allow functionality. For instance, the adult glomerular filtration barrier consists of fenestrated endothelium (50–100 nm diameter)(8), the glomerular basement membrane (300– 350 nm thick)(9), and podocyte foot processes (30–45 nm wide)(10), that together allow size- and charge-specific filtration. Variations to the dimensions, shape or composition of any of these features can lead to impaired glomerular filtration. Recently, glomerular ultrastructural features were shown to be predictive for clinical outcomes in proteinuric patients, confirming the powerful link between ultrastructure and functionality in the kidney.(11) Similarly, the cortical tubular epithelium is strictly polarized, forming continuous tubes with a mean diameter of 40 µm(12), equipped with different lengths of apical microvilli depending on the tubule type, increasing the surface area for transport and playing an essential role in mechanotransduction(13).

Clearly, the absence or change of such ultrastructural features alters kidney functionality and they therefore need to be faithfully replicated in kidney organoids. Eventually, the aim is that transplanted kidney organoids can restore kidney function *in vivo*, which requires the correct functional ultrastructure. Such recapitulation is also necessary for accurate disease modelling and drug testing.

In this manuscript, we compare the ultrastructure of kidney organoids and human fetal kidneys. From this analysis, we learned essential similarities and divergences in terms of the maturity of cell types and nephron segments, as well as features unusual to healthy fetal kidneys. We found an aberrant amount of glycogen throughout a variety of organoid cell types. We hypothesized that the culture is hyperglycemic. Accordingly, we found several features that are known for hyperglycemic cultures, such as tubular YAP activation, EGFR upregulation, mitochondrial damage, and tubular lipid accumulations. Compared to healthy human fetal kidneys, these features are unusual. These findings offer insight into the limitations of—and potential strategies to improve—kidney organoid culture.

## Methods

### Human fetal kidneys

The human embryonic and fetal material was provided by the Joint MRC/Wellcome Trust (grant# MR/R006237/1) Human Developmental Biology Resource (www.hdbr.org). The full details of the tissue are included in the Supplementary Methods and Supplementary Table 1. All fetal kidney data are published open access on the Image Data Resource (https://idr.openmicroscopy.org) under accession number idr0148.

### Induced pluripotent stem cell culture, differentiation and organoid culture

Induced pluripotent stem cells (iPSCs) were differentiated towards kidney tissue and the formed organoids were cultured as described in our previous work.(14) Briefly, the iPSC line LUMC0072iCTRL01 (Leiden University Medical Center, the Netherlands) was differentiated in APEL2 medium (STEMCELL Technologies) supplemented with 1% protein-free hybridoma medium (PFHMII, Thermo Fisher Scientific), 1% antibiotic-antimycotic (Thermo Fisher Scientific), 8 μM GSK-3 inhibitor CHIR99021 (R&D Systems), 200 ng/ml fibroblast growth factor 9 (FGF9; R&D Systems), and 1 μg/ml heparin (Sigma-Aldrich) over the course of 7 days (D7). The cells were then aggregated and further cultured as organoids on an air– liquid interface. From D7 + 5 onwards, growth factors and small molecules were stopped and the culture was continued in basic organoid culture medium (APEL2 with 1% PFHMII and 1% antibiotic-antimycotic). In total, five independent cultures were performed (N=5) unless specified otherwise. Full details of this culture are included in the Supplementary Methods.

### Embedding, processing and TEM imaging

The organoids were fixed and prepared for TEM imaging as described in the Supplementary Methods. Briefly, they were washed and subsequently stained with 1% osmium tetroxide and 1.5% K_4_Fe(CN)_6_ in 0.1 M sodium cacodylate. After washing with MilliQ and dehydrating in ethanol, the organoids were embedded in epon. The regions of interest (ROIs) were determined in toluidine blue–stained, semi-thin (1 µm) sections. Accordingly, ultrathin sections (60 nm) were cut, transferred to a 50 mesh copper grids and stained with 2% uranyl acetate in 50% ethanol and lead citrate.

Imaging was performed on a Tecnai T12 electron microscope equipped with an Eagle 4k × 4k CCD camera (Thermo Fisher Scientific). The ROIs chosen in toluidine blue sections were targeted in the TEM by making large scans with the EPU software (1.6.0.1340REL, FEI). Subsequently, the ROI was imaged in tiles at 2900× magnification and Omero (OME) and PathViewer were used to stitch, upload and annotate the data. For three independent cultures (N=3), one organoid per timepoint (n=1) was embedded and analyzed with TEM.

### Western immunoblotting

The Western immunoblot was performed as previously published.(14) Briefly, after snap-freezing in liquid nitrogen, the organoids were resuspended in 50–70 μL radioimmunoprecipitation assay lysis buffer (Sigma-Aldrich) supplemented with phosphatase inhibitor tablets (PhosSTOP, Sigma-Aldrich) and protease inhibitor (cOmplete ULTRA Tablets, Roche). The protein concentrations were determined using a Pierce BCA Protein Assay Kit (Thermo Fisher Scientific) according to the manufacturer’s protocol. Per well, 10 μg of protein was run on 10% acrylamide gels. All further steps were the same as previously published.(14) The protein bands were quantified by densitometry using ImageJ and normalized to GAPDH.(15) Primary and secondary antibodies used are listed in Supplementary Table 2. For five cultures (N=5), protein expression of one organoid per timepoint (n=1) was assessed. The results were analyzed in GraphPad Prism 9. After Shapiro-Wilk normality testing, an ordinary one-way ANOVA followed by a Tukey’s multiple comparisons test was performed.

### Immunofluorescence

The organoids were fixed at D7 + 5, D7 + 10, D7 + 14, D7 + 18, and D7 + 25 in 2% (v/v) paraformaldehyde (20 min, 4 °C). Cryopreservation and cutting, as well as immunofluorescence staining were performed as previously described (14) and are summarized in the Supplementary Methods. Briefly, after blocking and permeabilization, the cryo-sections were stained using indirect immunofluorescence with the antibodies listed in the Supplementary Table 2. After washing and mounting, the imaging was performed on an automated Nikon Eclipse Ti-E microscope equipped with a 20× air and a 40× oil objective. All images were processed using Fiji(16), without background removal or similar modifications. For five cultures (N=5), one organoid per timepoint (n=1) was assessed.

### Oil red O staining

Cryo-sections (15 µm) were made according to our previously published protocol.(14) The cryo-sections were equilibrated to RT and then washed with PBS warmed to 37 °C for 5 min to remove the embedding medium. After a quick rinse in 60% isopropanol in MilliQ, the sections were stained with Oil Red O (Sigma-Aldrich) for 20 min. Finally, after another rinse in 60% isopropanol, the slides were washed with MilliQ until it no longer turned red. The slides were mounted with Fluoroshield (Sigma-Aldrich) and imaged on an automated Nikon Eclipse Ti-E microscope using a 40× objective.

## Results

### Organoid glomeruli develop asynchronously and incorrectly compared to human fetal glomeruli

To determine the glomerular maturity in kidney organoids, their ultrastructure was compared to human fetal kidneys at multiple developmental stages. D7+18 organoid glomeruli were found to be comparable to several stages of fetal glomerular development, indicating asynchronous development compared to fetal kidneys. The overall macro-morphology of D7 + 18 organoid glomeruli was comparable to capillary loop stage 2 (CLS-2) in fetal kidneys (Figure 1a-b).

**Figure 1:**
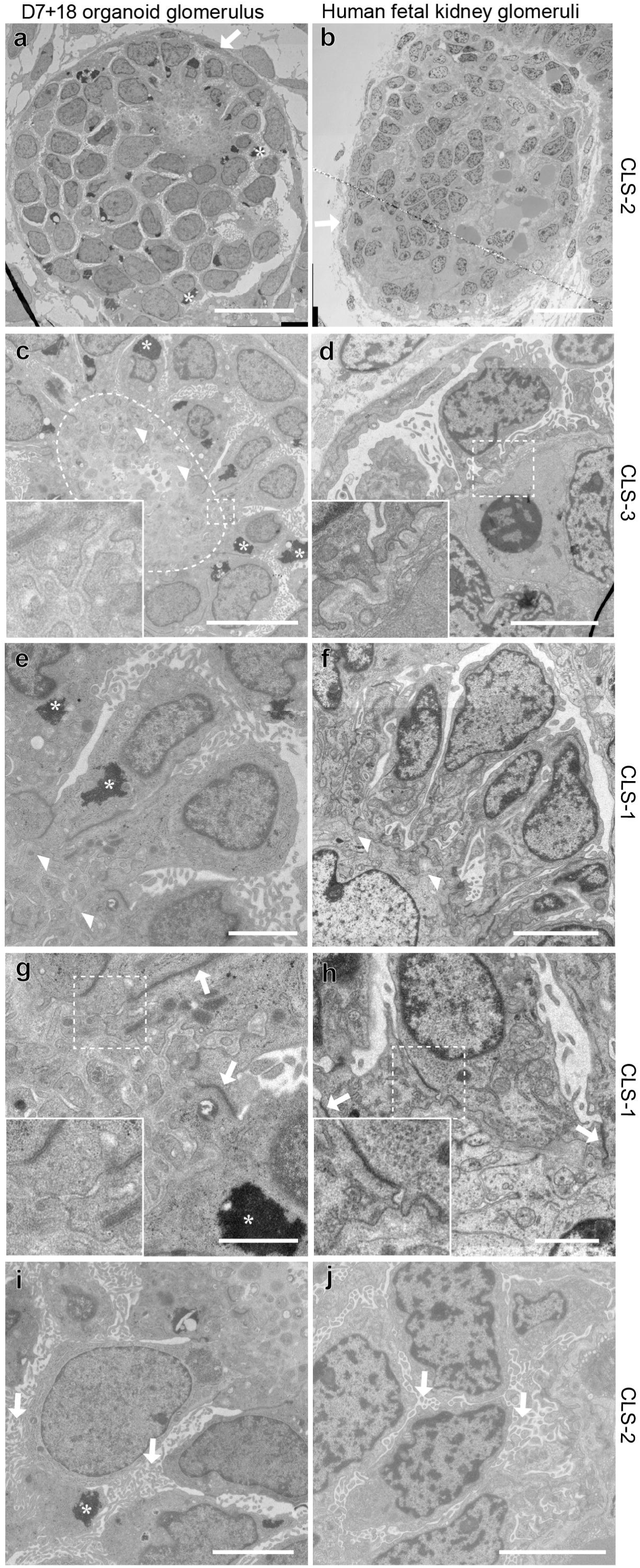
Congruent and divergent morphological features of glomerulogenesis in kidney organoids (left column) and human fetal kidneys (right column). (a) D7 + 18 organoid glomerulus with parietal-like cells (white arrow) had comparable macro-morphology to (b), a capillary loop stage (CLS)-2 glomerulus lined by parietal epithelial cells (PECs) (white arrow) and lack of a Bowman’s space. (c) Large areas of trilaminar basement membrane (inside dotted circle, zoom) were deposited in a disorganized manner, but in similar layers compared to (d) a CLS-3 stage glomerulus (zoom). (e) Podocytes that were attached to the basement membrane (white arrowheads) polarized similarly to (f) CLS-1 stage podocytes. (g) Large tight junctions (arrows) connected the apical side of polarized podocytes, which possessed primary foot processes comparable to (h) podocytes of the CLS-1. (i) Podocytes developed an excessive number of microvilli (arrows); more than at any stage in first trimester fetal kidneys, but most comparable to (j) CLS-2. Asterisks in all organoid panels label large glycogen deposits. All fetal kidney data were reproduced from our previously published open access library on the Image Data Resource (https://idr.openmicroscopy.org) under accession number idr0148. Patient IDs: b, d, j: 15333; f, h: 15335. Scale bars: a, b: 20 µm; c: 10 µm; d, f, i, j: 5 µm; e: 3 µm; g, h: 2 µm.

In both fetal kidneys and D7 + 18 organoids, a Bowman’s space was absent and morphologically a thin layer of parietal epithelial cells (PECs) surrounded the glomerulus (Figure 1a-b, arrow). However, immunofluorescence showed that cells in the D7 + 18 organoids expressed podocytes markers instead of PEC markers, indicating the absence of PECs (Figure S1).

Podocytes polarize upon glomerular basement membrane (GBM) deposition by endothelial cells and podocytes.(7) Notably, in kidney organoid glomeruli, the endothelial cells were absent. Instead, large areas of irregularly shaped basement membrane accumulated in various locations throughout the glomeruli (Figure 1c, inside dotted circle) and consequently, podocytes at D7 + 18 did not polarize homogenously (Figure 1a). Only podocytes directly adjacent to the areas of the GBM adhered with early foot processes (Figure 1c, e, arrowheads), similar to CLS-1 podocytes (Figure 1f, white arrowheads). Podocytes beyond those areas lacked polarization. Interestingly, although irregularly shaped, the GBM was composed of the three layers found in CLS-3 glomeruli (lamina densa (dark gray), lamina rara externa and lamina rara interna (light gray), Figure 1d), which also exist in mature GBM *in vivo*(17). At the same time, podocytes were connected to each other via tight junctions both at the cell body and apical side (Figure 1g, arrows), as seen in the CLS-1 stage (Figure 1h, arrows). Furthermore, podocytes possessed microvilli at D7 + 18 (Figure 1i, arrows), however, these were excessively abundant compared to fetal kidneys (Figure 1j, arrows). Prominent intracellular glycogen deposits were also observed (Figure 1c, e, g, i, asterisks), which were not seen in glomeruli in fetal kidneys of the first trimester (Figure 1b).

### Kidney organoid cells show aberrant glycogen deposits along with mitochondrial damage

Glycogen deposits were abundant in various cell types in kidney organoids, unlike in healthy fetal kidneys. At D7 + 25 in the kidney organoids, glycogen accumulated predominantly in podocytes (Figure 1 a,c,e,g,i, white asterisks) and in chondrocytes within glomeruli, and to a lesser extent in tubules and interstitial cells (Figure S2). In contrast, glycogen was absent in glomerular cells, tubular cells, and the interstitium in fetal kidneys (Figure 2a-f), where only the collecting duct possessed large deposits (Figure 2g).

**Figure 2:**
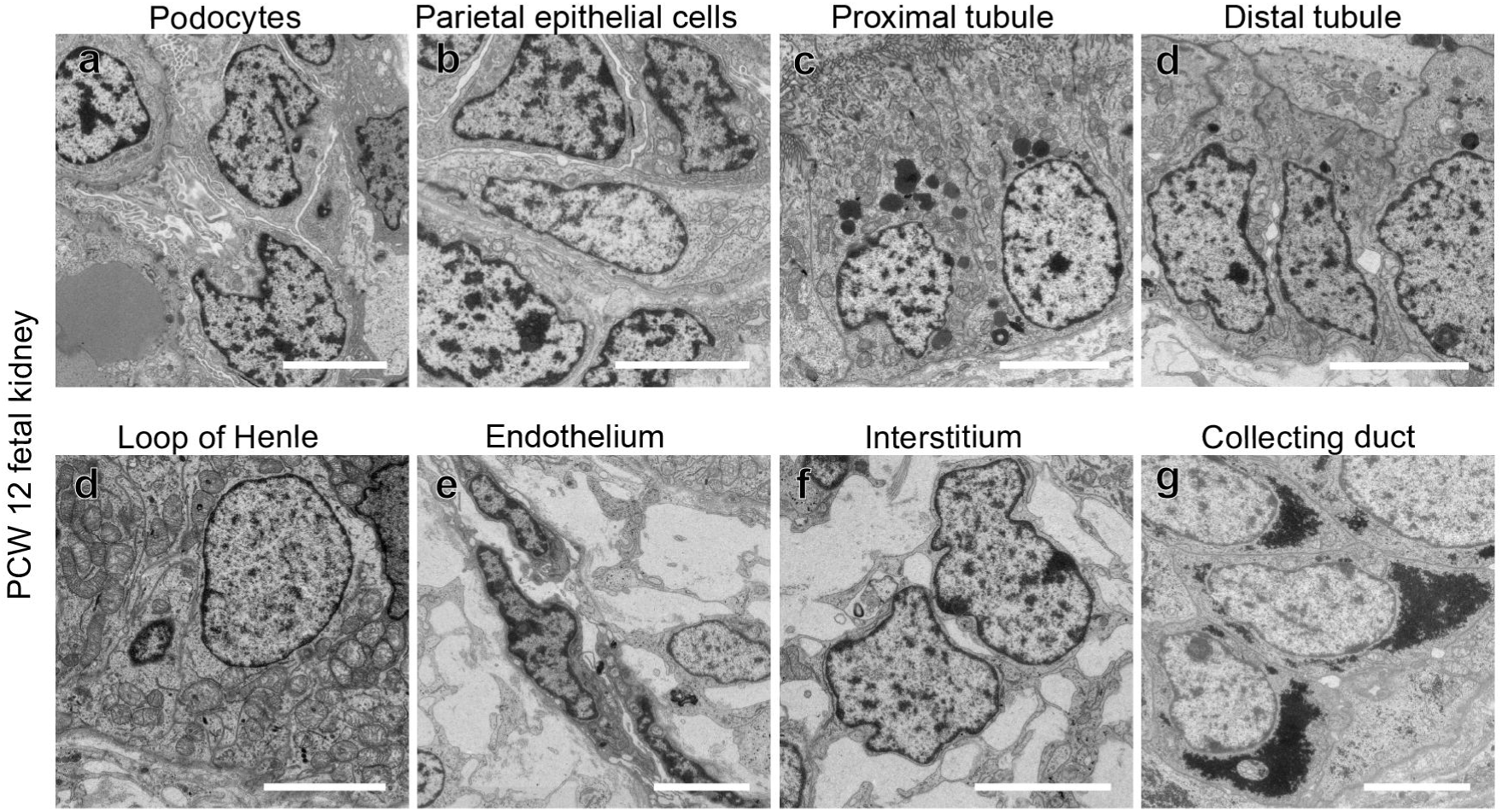
Glycogen deposits in week 12 fetal kidneys. (a-f) Different renal cell types do not possess large intracellular glycogen deposits, except for (g) collecting duct cells (arrows). Patient ID: a-b: 15317; c-g: 15335; scale bars: 5 µm. Post-conceptional week (PCW).

Given their prominence, our next aim was to understand if the glycogen deposits were present throughout the entire culture protocol and how they potentially associated with cell type–specific ultrastructural features. For this, D7 + 5, D7 + 10, D7 + 14, D7 + 18 and D7 + 25 organoids were analyzed. At D7 + 5, we did not detect glycogen deposits in toluidine blue stained semi-thin sections, but we found them in abundance beginning at D7 + 10 (Figure S3). The first glycogen deposits appeared surrounding lipid vesicles at D7 + 10, predominantly in stromal cells and rarely in podocytes (Figure 3a-c, Figure S3b). Furthermore, we noticed mitochondrial abnormalities, characterized by a decrease of cristae at D7 + 10 compared to D7 + 5 and an increase in electron-lucent areas (Figure 3c). Ribosome-covered rough endoplasmic reticula (ER) were directly adjacent to the mitochondria (Figure 3c, inset). Later in culture, at D7 + 18, the amount of glycogen increased significantly (Figure 2d-e) and mitochondrial damage persisted (Figure 3f). At the last time point in culture, D7 + 25, glycogen deposits were comparable to D7 + 18; however, an increased amount of cell debris as well as the appearance of glycogen-containing and ECM-producing chondrocytes adjacent to podocytes was noticed (Figure 3g-h).

**Figure 3:**
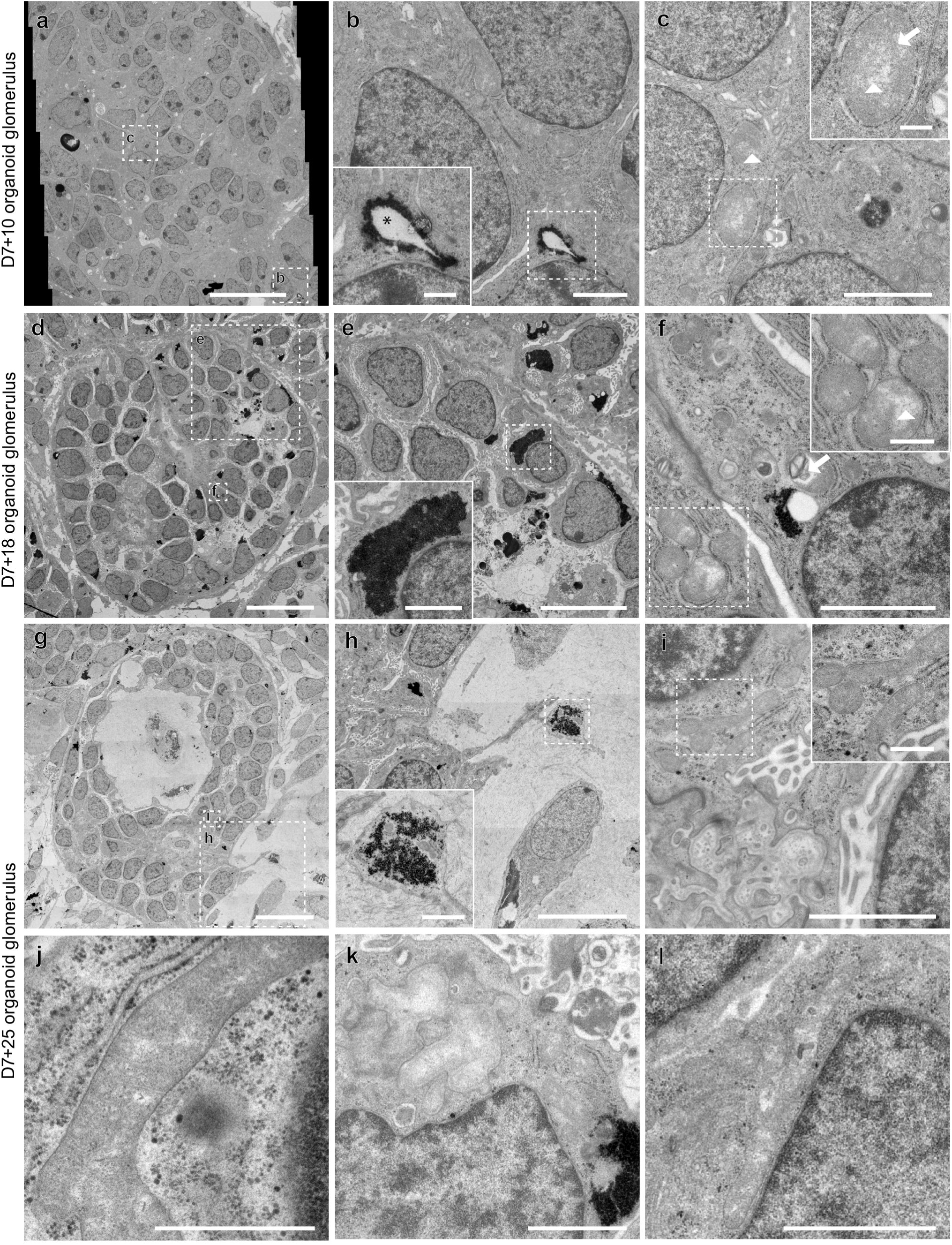
Glycogen deposits and mitochondrial abnormalities in kidney organoid glomeruli increased over time. In (a) glomeruli at D7 + 10, glycogen deposits were rare. (b) Glycogen deposits surround lipid vesicles. (c) Remnants of mitochondrial cristae were visible in podocytes (arrow) and mitochondria contained large electron-lucent areas (arrowheads). (d) Glomeruli at D7 + 18 contain an increased amount of intracellular glycogen and (e) the size of the glycogen deposits had increased. (f) Aberrant mitochondrial morphology persisted. (g) Glomeruli at D7 + 25 contained chondrocytes producing large areas of ECM. (h) Intracellular glycogen deposits were located in podocytes and chondrocytes. (i-l) Mitochondrial morphology was diverse, but the scarcity of cristae and the presence of electron-lucent areas were uniform. Scale bars: a, d, g = 20 µm; e, h = 10 µm; c = 3 µm; b, e (zoom), f, h (zoom), i, k, l = 2 µm; b (zoom), c (zoom), f (zoom), i (zoom), j = 0.6 µm.

Mitochondrial morphology was diverse at D7 + 25. While the minority of cells had mitochondria with cristae (Figure 3i, j), occasionally mitochondria were swollen (Figure 3k) and more frequently fused (Figure 3j,l). The majority of podocyte mitochondria possessed electron-lucent areas (Figure 3j-l). Overall, we noticed an increase in glycogen deposits in a variety of cell types over time along with the deterioration of mitochondrial morphology.

### Glycogen synthase kinase 3 beta (GSK3β) phosphorylation increased over time

The prominent glycogen deposits in the organoids indicated an aberrant metabolism compared to fetal kidneys. Glycogen synthase kinase 3 (GSK3), a constitutively expressed multi-functional serine threonine kinase, negatively regulates glycogen synthesis both *in vitro*(18) and *in vivo*(19). We therefore sought to determine its activity in the organoid culture and expected an increased inactivation over time, comparing D7 + 5, D7 + 10, D7 + 14, D7 + 18 and D7 + 25. Protein expression was analyzed by Western blot, which showed a significant increase (*p* < 0.0001) of phosphorylated GSK3β over time in culture (Figure 4a,d, Figure S4a-e). Total GSK3β was unchanged (Figure 4b,e), therefore the ratio of phosphorylated to total GSK3β also significantly increased (*p* < 0.0001) (Figure 4c, Figure S4f-j).

**Figure 4:**
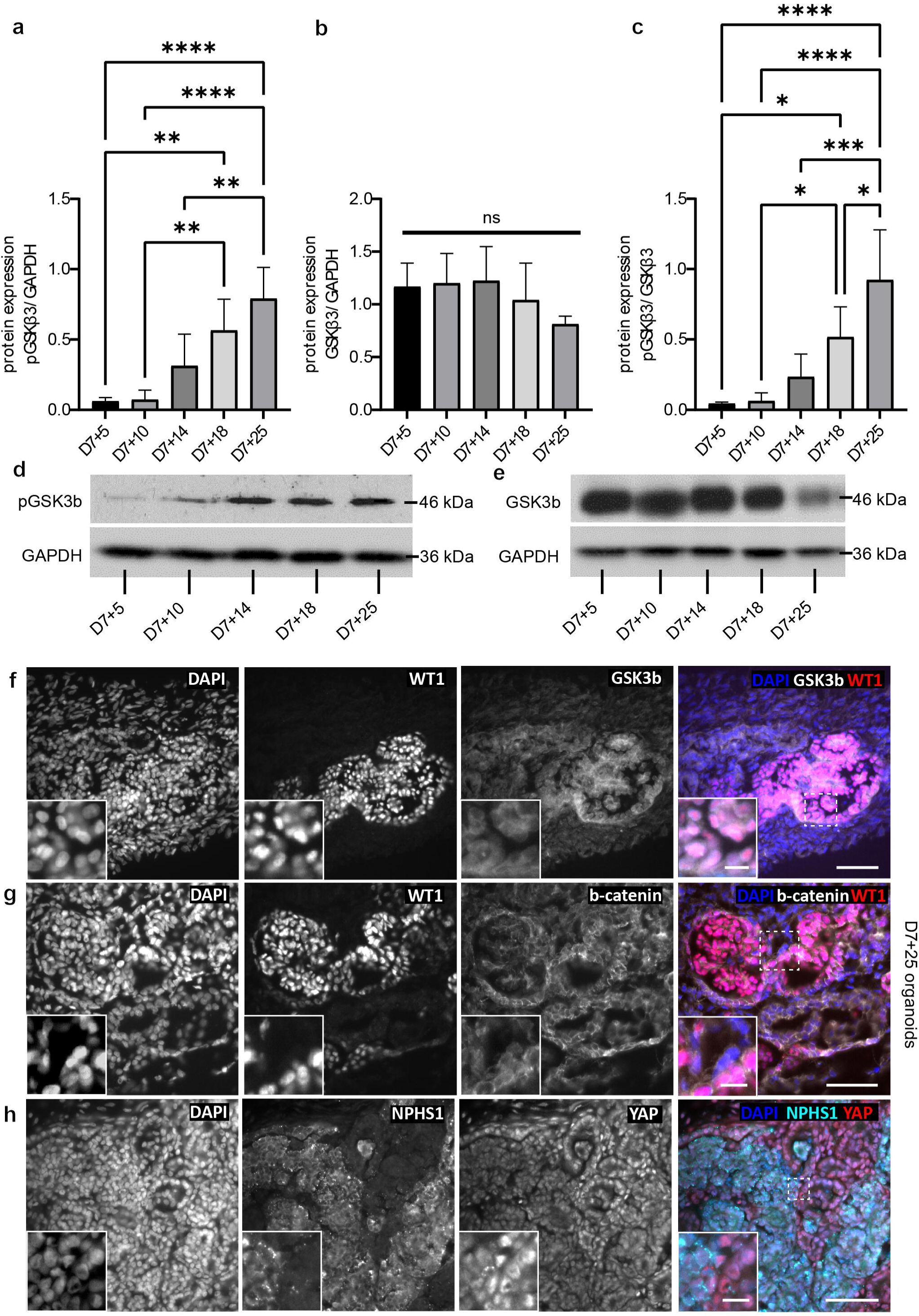
pGSK3β, β-catenin and YAP expression in kidney organoids. (a,d) GSK3β was increasingly phosphorylated in kidney organoids over time. (b,e) Total GSK3β expression remained unchanged. (c) The ratio of total GSK3β and pGSK3β increased over time in culture. (f) GSK3β was expressed mainly in WT1^+^ podocytes at D7 + 25. (g) β-catenin was expressed in its inactive form, as seen by the cytoplasmic expression in WT1^+^ podocytes and tubules at D7 + 25. (h) YAP existed in both an inactive and an active form, as seen by its cytoplasmic and nuclear expression in NPHS1^+^ podocytes and nuclear expression in tubules at D7 + 25. Scale bars: 50 µm, insets: 10 µm. *: p < 0.05, **: p < 0.005, ***: p = 0.0003, ****: p < 0.0001.

### Aberrant YAP activation in proximal tubular cells

GSK3 inhibition in podocytes is known to lead to their apoptosis, and is regulated by the Hippo pathway independent of Wnt-β-catenin signaling.(20) This incentivized us to determine the expression and localization of GSK3β (an essential component of the β-catenin destruction complex(21)), β-catenin (a key component of the Wnt signaling pathway and the substrate of GSK3(21, 22)) and YAP (a downstream effector of the Hippo pathway that likewise associates with the β-catenin destruction complex, co-regulating proliferation and apoptosis(23, 24)). Immunofluorescence staining of D7 + 25 organoids showed that total GSK3β localized predominantly to podocytes, but also to tubular cells (Figure 4d). In the absence of Wnt, β-catenin is known to be ubiquitinated by the GSK3-containing destruction complex, and consequently degraded.(23) In the organoids, β-catenin expression was cytoplasmic (Figure 4e), indicating the absence of canonical Wnt signaling. Similar to β-catenin, YAP can be degraded by the destruction complex involving GSK3. However, in contrast to β-catenin, YAP expression was both nuclear and cytoplasmic in podocytes (Figure 4f) with no clear difference in expression over time (between D7 + 10 and D7 + 25) (Figure S5). Furthermore, nuclear YAP was more highly expressed in tubules compared to podocytes at D7 + 25 (Figure 4f). Particularly LTL^+^ proximal tubules strongly expressed nuclear YAP (Figure 4f, Figure S5). While nuclear YAP in podocytes is critical for podocyte survival *in vitro* and *in vivo* (25, 26), YAP activation in proximal tubules is associated with diabetic kidney disease and high glucose *in vitro* cultures (27, 28). We therefore investigated additional hallmarks of hyperglycemia and diabetic kidneys, such as epidermal growth factor receptor (EGFR) upregulation and activation, lipid accumulations, and mitochondrial damage.

### EGFR was increasingly upregulated and activated

YAP is known to be positively regulated by EGFR, and YAP activation therefore indicates active EGFR signaling. Although EGFR is necessary in nephrogenesis, predominantly in the ureteric bud(29), its upregulation and activation can be pathological and are associated with hyperglycemia and diabetic nephropathy.(30) In the organoids, protein levels of both pEGFR (*p* < 0.0042) and EGFR (*p* < 0.0008) significantly increased over time from D7 + 5, D7 + 10, D7 + 14, D7 + 18 to D7 + 25 (Figure S6). This is in line with the activated YAP found in proximal tubules.

### Glycogen and lipid droplets accumulated in tubules with damaged mitochondria

Since tubules contained nuclear YAP expression, which is associated with tubular damage, we were interested to morphologically assess tubular development in the organoids. Accordingly, we found increasing glycogen deposits and lipid droplets over time in tubules and the mitochondria showed morphological aberrations. At D7 + 10, tubules rarely contained glycogen and the cristae in mitochondria were present, but dim (Figure 3a-c). At D7 + 18, tubules contained intracellular lipid droplets (Figure 5d-e). Dim cristae persisted in mitochondria and electron-lucent areas developed (Figure 3f), associated with damage to the mitochondrial matrix(31–33). At D7 + 25, glycogen and lipid accumulations were mutually exclusive in tubules. Tubules at D7 + 25 contained either an excess of lipid vesicles (Figure 5g-i) or glycogen deposits (Figure 5j-l). In either type of tubule, cristae were difficult to distinguish in mitochondria, and electron-lucent areas were present (Figure 5i,k), similar to D7 + 18. The lipid droplets had an unusual morphology (Figure 5d-i). Unlike their well-known definition, they had a clearly distinguishable membrane and were heterogenous, containing electron-dense material (Figure 5i, arrowheads). To confirm them to be lipid droplets, we stained cryo-sections with Oil red O, a reagent that binds neutral triglycerides and lipids (Figure 6). At D7 + 10, small lipid droplets were present throughout the organoids and partially localized to tubules (Figure 6a). At D7 + 25, the number and size of lipid droplets increased and were mainly present in tubules (Figure 6b). Small droplets could also be noticed in locations such as in chondrocytes, in which glycogen deposits with lipid droplets were seen in TEM (Figure S3, arrowhead). Overall, the described features could not be assigned to a tubule type, since discrimination of proximal and distal tubules in the organoids based on ultrastructure or histology was impossible. The tubules were morphologically homogenous and were not comparable to the diversity in human fetal kidney tubules(7).

**Figure 5:**
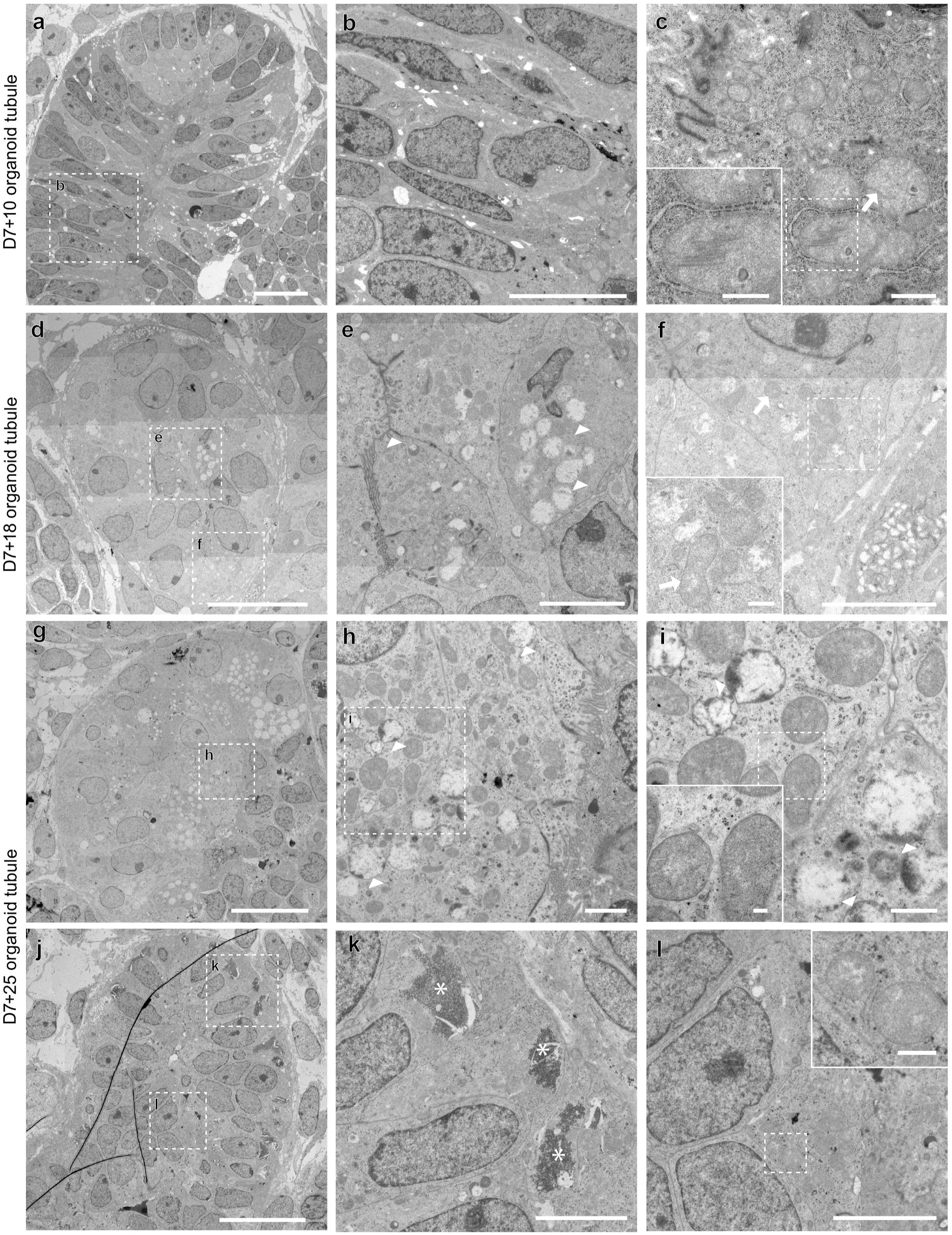
Intracellular accumulations of glycogen and lipid droplets in kidney organoid tubules over time. (a-b) D7 + 5 tubules did not contain glycogen deposits or lipid droplets and (c) cristae in mitochondria were dim, but visible (arrows). (d-e) D7 + 18 tubules contained a small number of lipid droplets (arrowheads). (f) Cristae were less visible (arrow), and mitochondria developed electron-lucent areas. At D7 + 25, tubules contained either (g-i) large numbers of lipid droplets (arrowheads) or (j-l) large accumulations of glycogen (asterisks) and the mitochondrial morphology was comparable to D7 + 18. Scale bars: a,d,g,j: 20 µm; b: 10 µm; e,f,k,l: 5 µm; h: 2 µm; c,c (zoom),i: 1 µm; f (zoom),l (zoom): 0.6 µm; i (zoom): 0.2 µm.

**Figure 6:**
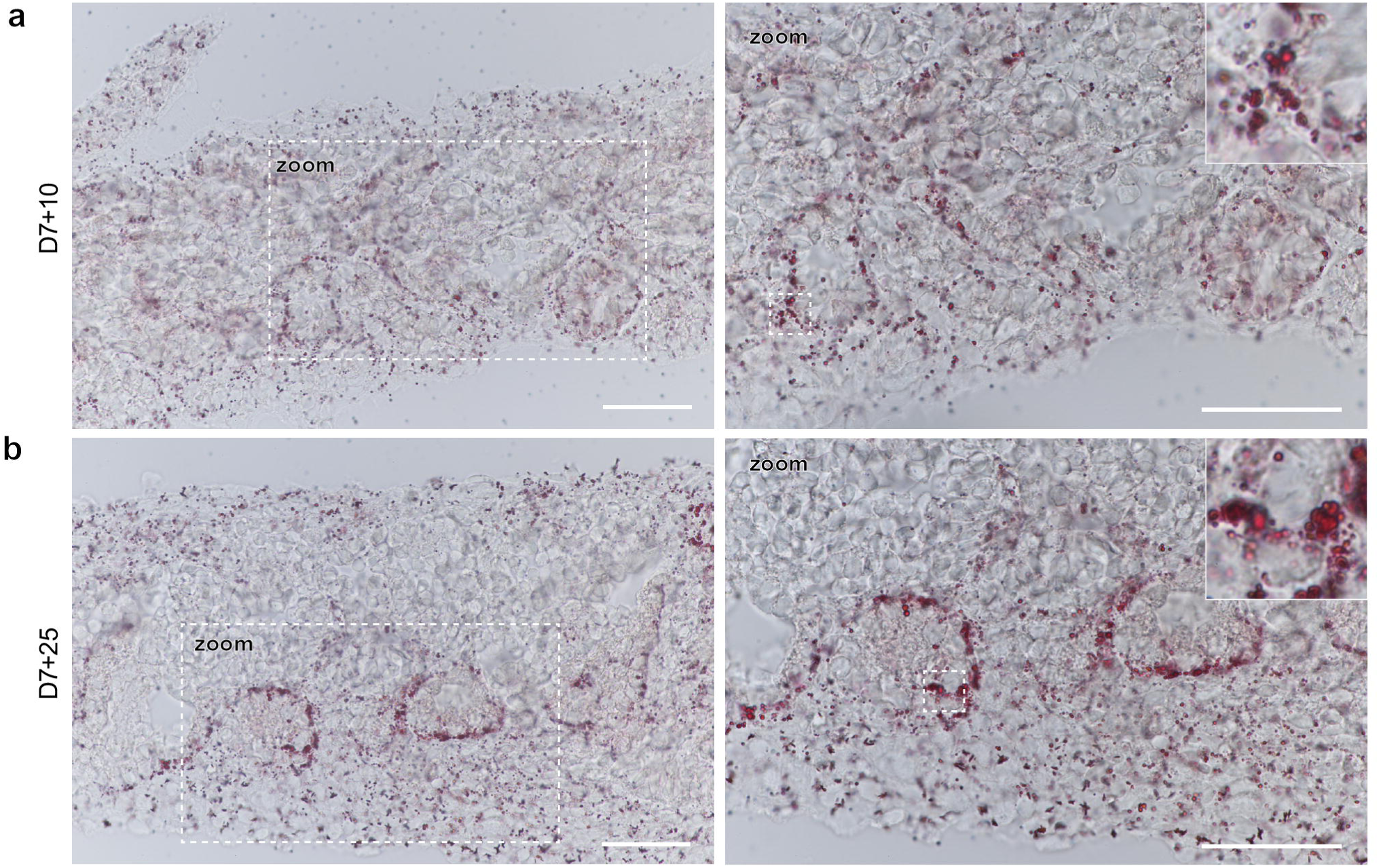
Oil red O staining of lipid droplets in D7 + 10 and D7 + 25 organoids. (a) Lipid droplets were present throughout D7 + 10 organoids and partially localized to tubules. (b) In D7 + 25 organoids, the number and size of lipid droplets increased compared to D7 + 10 organoids and mainly localized to tubules. Dashed box: zoomed images in right column. Scale bars: 50 µm.

## Discussion

Whether destined for future transplantation or for disease modeling *in vitro*, it is essential that kidney organoids correctly replicate the functional ultrastructure of the kidney. Therefore, the initial aim of the study was to assess morphological differences in kidney organoids compared to human fetal kidneys using TEM. On top of differences in nephron morphology, we noticed features associated with hyperglycemic cultures and diabetic nephropathy.

Initially, we noted large intracellular glycogen deposits, which became increasingly abundant over time. Yet, glycogen accumulation is unusual compared to human fetal kidneys outside the collecting duct (Figure 2).(7) Morphologically, the glycogen deposits in our culture were similar to deposits described by a study that inactivated GSK3 using the Wnt activator CHIR99021 in pluripotent stem cells (18), which made us wonder if the CHIR99021 treatment early in our culture protocol could be the cause. However, the significant increase in phosphorylated GSK3β over 20 days of organoid culture (Figure 4a-c) and the low pGSK3β expression at D7 + 5 (two days after the last CHIR99021 treatment) indicated a different underlying cause.

The accumulation of glycogen induced by GSK3b inhibition is just one of several features we found in our organoid culture that are known to coincide in diabetic nephropathy and hyperglycemic cultures. GSK3 inhibition has recently been studied in the context of podocyte health.(20) Accordingly, upon inactivation or absence, podocytes express activated YAP and re-enter the cell cycle until they reach mitotic catastrophe. Cell cycle re-entry of mature podocytes induced by GSK3 inhibition, coinciding with YAP activation, is an early hallmark of diabetic nephropathy.(34) Additionally, hyperglycemia is known to lead to intracellular accumulation of glycogen in diabetic kidneys.(35) Similar to these studies, GSK3β was increasingly phosphorylated in our organoids (Figure 4a) and Wnt was inactive (cytoplasmic β-catenin expression, Figure 4e). YAP was expressed in both the nucleus and cytoplasm, even late in culture (D7 + 25), which is known to be essential in podocytes of both developing and adult kidneys.(26, 36) Yet, in combination with GSK3b inhibition and glycogen accumulation, it could indicate a hyperglycemic culture.

Similar to podocytes, tubules showed features of hyperglycemia and diabetic nephropathy. Nuclear (activated) YAP was strongly expressed in proximal tubules (Figure 4f, Figure S5), which is known to drive tubulointerstitial fibrosis in diabetic nephropathy.(27) This is in line with the fibrotic interstitial phenotype we found in our earlier study.(37) Furthermore, EGFR signaling significantly increased over time in our kidney organoid culture (Figure S6). EGFR upregulation and activation are regulated by serine/threonine protein kinase (SKG-1), which in turn is upregulated by TGF-β1, a process essential for the onset of diabetic nephropathy and well-known for hyperglycemic cultures.(38) Knockdown of EGFR expression in hyperglycemic cultures co-reduces YAP expression and phosphorylation, confirming the link between EGFR and YAP.(28) EGFR signaling also increased in podocytes when cultured in high glucose.(39) Overall, there is a well-known link between EGFR and YAP activation in renal cells in hyperglycemia and diabetic nephropathy, and our data suggest that the same link is present in kidney organoids.

In line with our hypothesis of a hyperglycemic environment, we found excessive lipid accumulation in tubules of the kidney organoid culture that increased over time (Figure 5, 6). In diabetic kidneys and hyperglycemic cultures *in vitro*, beta oxidation is decreased, decreasing ATP production and leading to lipid accumulation. This has been mainly described in tubules, and is frequently referred to as lipotoxicity in humans and rodents, as well as *in vitro*.(34, 40–47) Furthermore, phosphorylation of GSK3β, as seen in the organoids, is known to be involved in high-glucose mediated lipid depositions in renal tubular cells.(47) A recent study of this kidney organoid model showed that, with the absence of CPT1, the proximal tubules fail to use the mitochondrial long-chain fatty acid metabolism (48), confirming our findings. The altered metabolism induced by high glucose also leads to mitochondrial damage due to the production of reactive oxygen species (ROS). Mitochondria are the major site of ROS production (49) and therefore mitochondrial damage and dysfunction as a result of hyperglycemia have been frequently described *in vitro*(50, 51) and in a variety of tissues *in vivo*, including the kidney(52–58). In our study, the majority of mitochondria in kidney organoids rarely or never contained cristae and instead showed electron-lucent areas. A comparable morphology has been shown, for instance, in the mitochondria of an insulin-resistant rat model(49), in endothelial cells and podocytes treated with diabetic serum(59), and in hyperglycemia in diabetic kidneys along with mitochondrial hyperbranching and fragmentation/fission(40, 60). For these reasons, we hypothesized that the standard organoid culture medium might be hyperglycemic.

In line with our hypothesis, measurements showed that the organoid culture indeed was hyperglycemic. The glucose concentration of the kidney organoid culture medium (2.92 g/L, Figure S7) was non-physiological compared to basal plasma concentrations (0.7–1.0 g/L, (61, 62)) and represents a pre-diabetic milieu.(63) Cell culture media usually range from 1–10 g/L glucose, whereby 1.8 g/L is already considered supraphysiological and a pre-diabetic milieu and 5 g/L and higher are used for diabetic models. A recent kidney organoid study measured a similar concentration(48) and the same organoid culture medium has also been used in another kidney organoid study where glycogen deposits can be seen in TEM images(64), confirming our findings.

Relating these findings back to the morphological development of the kidney organoids, the hyperglycemic environment might be one explanation for the insufficient vascularization of the organoids and particularly for the absence of endothelium in glomeruli. *In vitro*, serum derived from diabetic mice induced mitochondrial dysfunction, and significantly reduced NOS activity leading to endothelial dysfunction and secretion of proapoptotic factors inducing podocyte apoptosis.(59) *In vivo*, similar to proximal tubules, YAP is activated in endothelial cells in diabetic kidneys(65) and inhibition reduced inflammatory markers and monocyte attachment. Since endothelial cells together with podocytes build the GBM early in nephrogenesis(7, 66), the absence of endothelial cells certainly influences the architecture of glomeruli and can explain the differences with fetal kidneys we found over the course of glomerular maturation (Figure 1). The excess of microvilli on podocytes is reminiscent of microvillus transformation as seen in podocyte injury(67–69), a feature that has had little attention in diabetic nephropathy.(70) Late-stage diabetic nephropathy/hyperglycemic features such as basement membrane thickening and podocyte detachment (71) are challenging to define in this culture, due to the heterogeneous morphology of basement membrane and podocytes as well as the emergence of chondrocytes. Though, the latter is a possible explanation for the excessive matrix depositions within the organoids (Figure 1c, 3g-h).

The excessive matrix depositions likely derive from the off-target chondrocyte population. GSK3 inhibition is known to induce maturation of chondrocyte progenitors (72, 73), which is in line with the glycogen deposits in and abundance of matrix around the chondrocytes in the kidney organoids (Figure 3g-h). The chondrocyte population potentially emerges from mesangial cells, which in diabetic nephropathy obtain a chondrogenic phenotype.(74) Furthermore, induced by the activation of EGFR, mesangial cells increasingly produce collagen type 1 in diabetic nephropathy (75, 76), which is one of the ECM proteins that was strongly upregulated in the stromal population in our previous study.(37)

In summary, when comparing kidney organoid morphology to human fetal kidneys, we encountered excessive accumulations of glycogen, along with an increase in GSK3β phosphorylation, YAP activation in proximal tubules, and upregulated EGFR signaling. Furthermore, mitochondrial morphology indicated injury and excessive lipid accumulations were found in tubules. These findings are well-known for hyperglycemic cultures and diabetic nephropathy. Finally, from a different perspective, we believe this kidney organoid model could potentially be a suitable model for diabetic nephropathy. Future studies could further investigate the metabolic and morphological changes to find therapeutics for disease prevention or progression.

## Disclosure statement

The authors declare no competing interests.

## Data sharing statement

All fetal kidney data were reproduced from our previously published open access library on the Image Data Resource (https://idr.openmicroscopy.org) under accession number idr0148. All data generated for this article was made publicly available for download from https://doi.org/10.34894/BNDNTW.

## Funding source

This research did not receive any specific grant from funding agencies in the public, commercial, or not-for-profit sectors.

## Acknowledgements

The authors wish to thank Dr Steven Lisgo and Berta Crespo Lopez from the HDBR for their high-quality and quick work in providing the fetal kidney tissue for this study. Furthermore, the authors would like to thank the iPSC facility of the LUMC for the iPSC cell line and Prof. Wilfred Germeraard and his team from CiMaas for determining the glucose concentration of the organoid culture medium. Finally, we would like to thank the Microscopy CORE Lab members of Maastricht University for their scientific and technical support.

## Notes

https://doi.org/10.34894/BNDNTW

### Competing Interest Statement

The authors have declared no competing interest.

