## Supplementary Figures for "Ultrastructural comparison of human kidney organoids and human fetal kidneys reveals features of hyperglycemic culture"

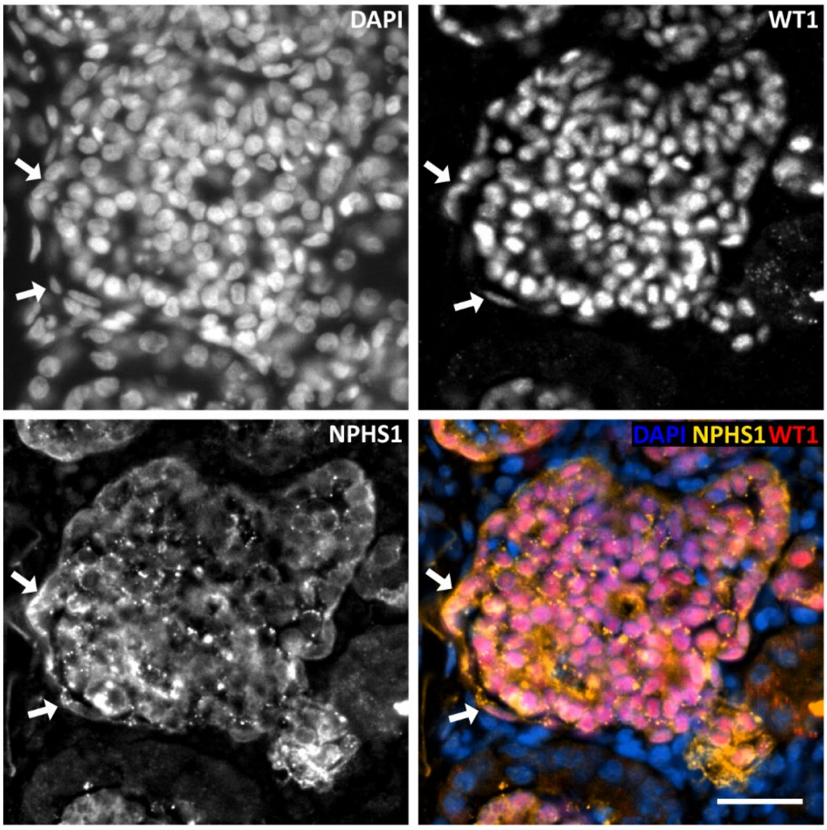


*Figure S1: Immunofluorescence staining of D7 + 25 cryosections of kidney organoids. The glomeruli are stained with the podocyte markers Nephrin (NPHS1) and Wilm's Tumor protein 1 (WT1), showing that elongated cells surrounding the glomerulus express podocyte markers (arrows). Scale bar: 25 µm.*

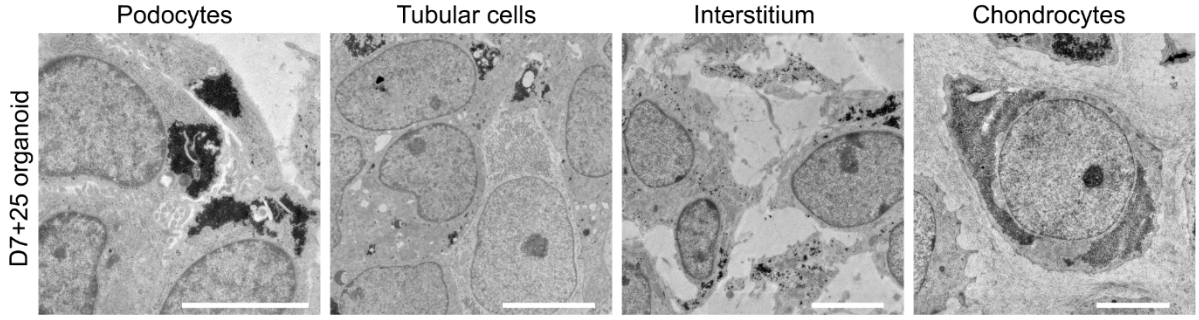


*Figure S2: TEM images showing glycogen deposits (black granules) in D7 + 25 kidney organoids. Scale bars: 5 µm.*

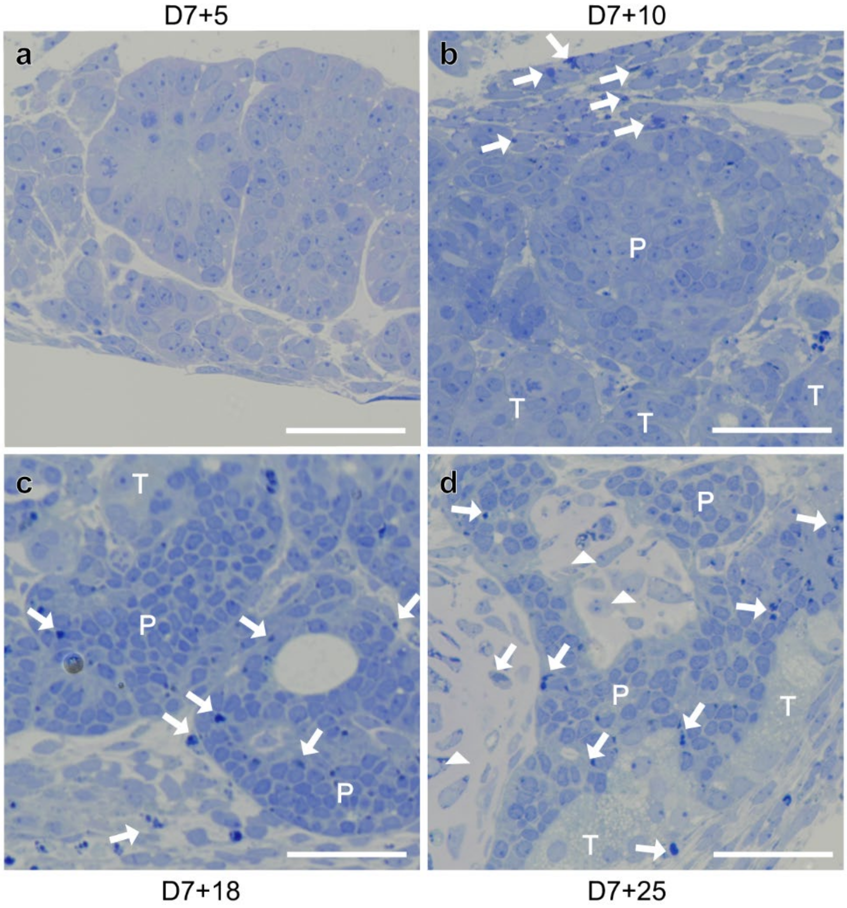


*Figure S3: Toluidine blue stained sections of kidney organoids show increased accumulation of glycogen (arrows) throughout the organoid over time and appearance of glycogen containing cartilage (arrowheads). Scale bars: 50 µm. T = tubular cells, P= podocytes.*

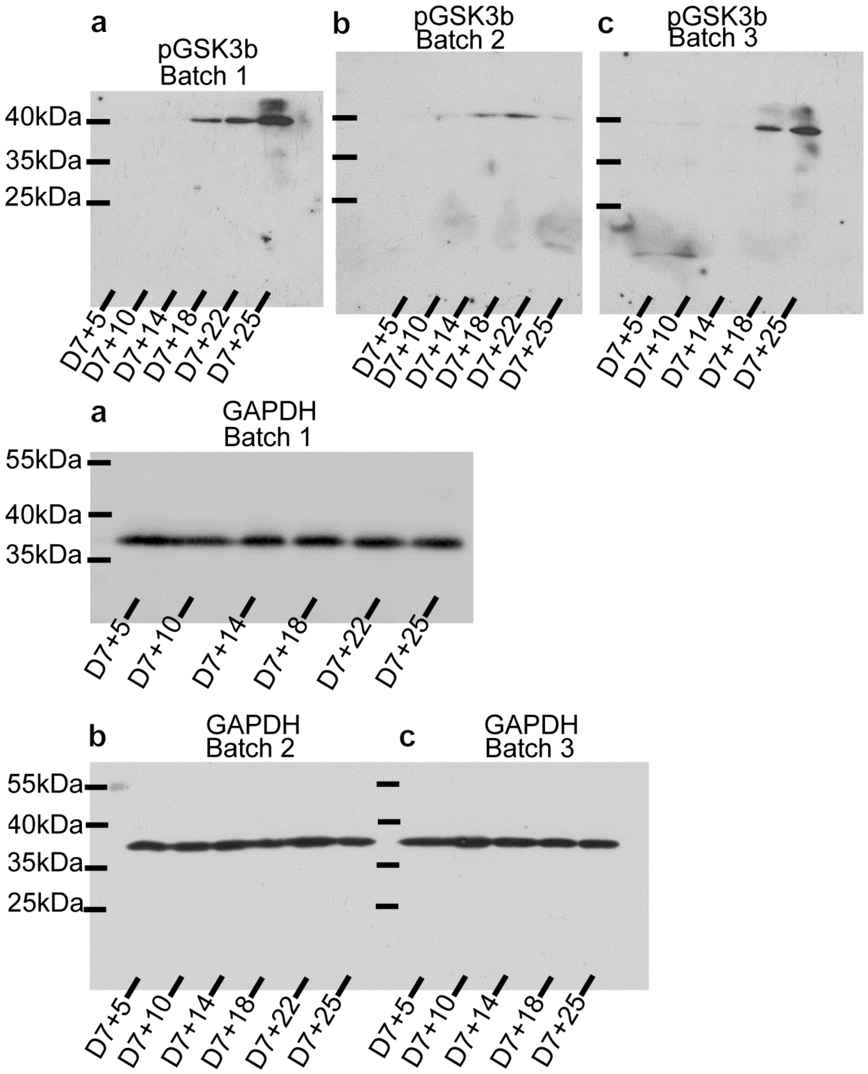
a
pGSK3b
Batch 1
40kDa
35kDa
25kDa
D7+5
D7+10
D7+14
D7+18
D7+22
D7+25
b
pGSK3b
Batch 2
D7+5
D7+10
D7+14
D7+18
D7+22
D7+25
c
pGSK3b
Batch 3
D7+5
D7+10
D7+14
D7+18
D7+25
a
GAPDH
Batch 1
55kDa
40kDa
35kDa
D7+5
D7+10
D7+14
D7+18
D7+22
D7+25
b
GAPDH
Batch 2
55kDa
40kDa
35kDa
25kDa
D7+5
D7+10
D7+14
D7+18
D7+22
D7+25
c
GAPDH
Batch 3
D7+5
D7+10
D7+14
D7+18
D7+25

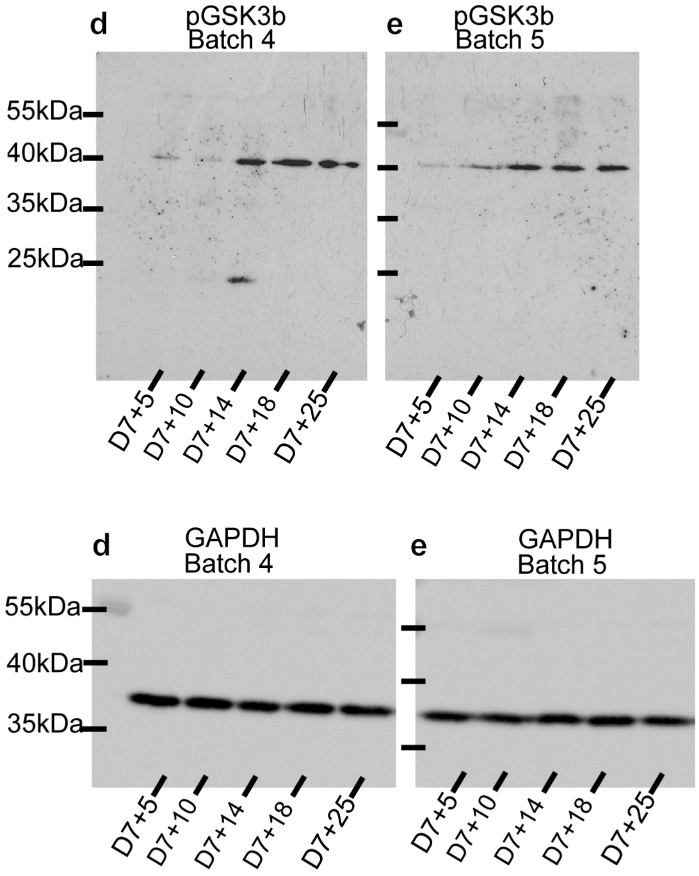
d
pGSK3b
Batch 4
e
pGSK3b
Batch 5
55kDa
40kDa
35kDa
25kDa
D7+5
D7+10
D7+14
D7+18
D7+25
D7+5
D7+10
D7+14
D7+18
D7+25
d
GAPDH
Batch 4
e
GAPDH
Batch 5
55kDa
40kDa
35kDa
D7+5
D7+10
D7+14
D7+18
D7+25
D7+5
D7+10
D7+14
D7+18
D7+25

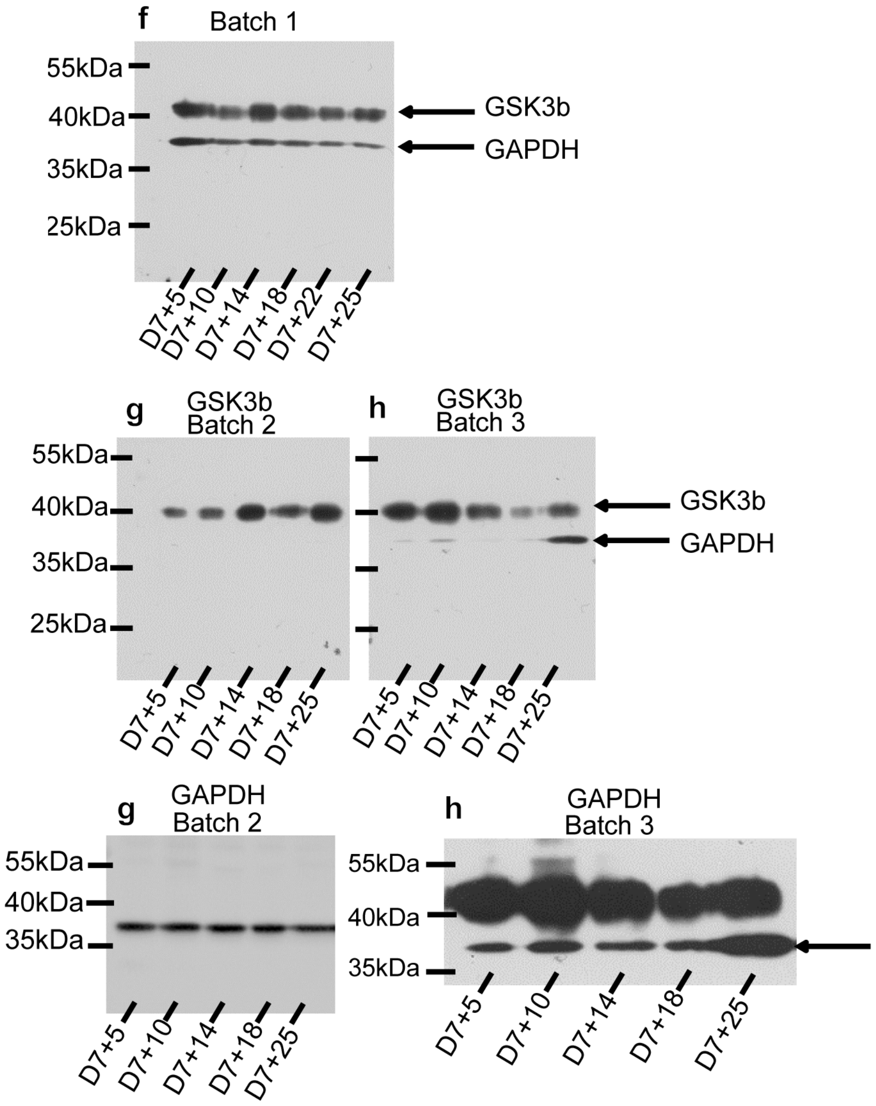
f
Batch 1
55kDa
40kDa
35kDa
25kDa
GSK3b
GAPDH
D7+5
D7+10
D7+14
D7+18
D7+22
D7+25
g
GSK3b
Batch 2
h
GSK3b
Batch 3
55kDa
40kDa
35kDa
25kDa
GSK3b
GAPDH
D7+5
D7+10
D7+14
D7+18
D7+25
D7+5
D7+10
D7+14
D7+18
D7+25
g
GAPDH
Batch 2
55kDa
40kDa
35kDa
D7+5
D7+10
D7+14
D7+18
D7+25
h
GAPDH
Batch 3
55kDa
40kDa
35kDa
D7+5
D7+10
D7+14
D7+18
D7+25

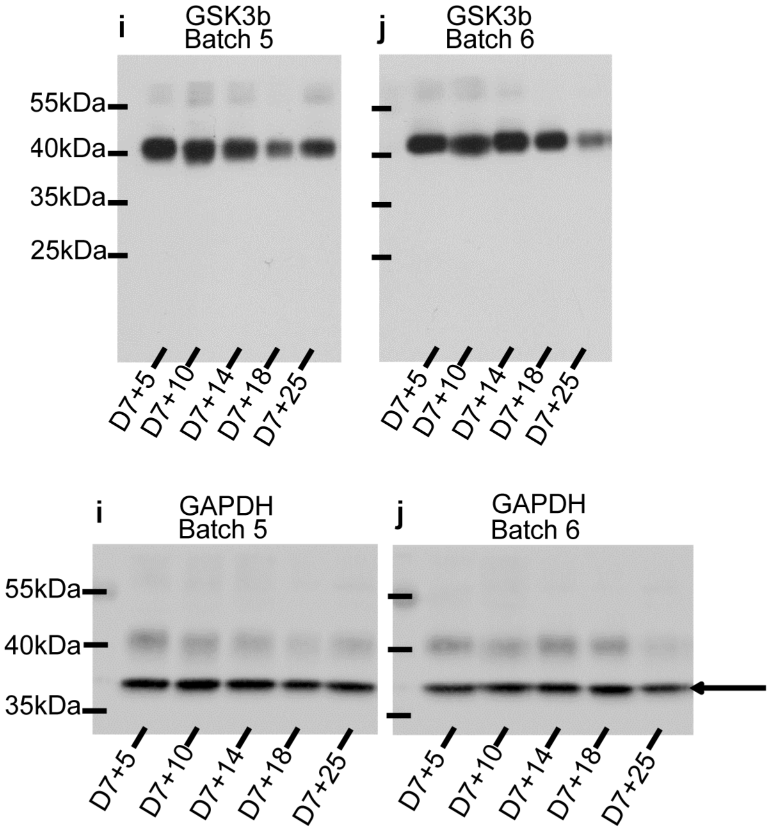


*Figure S4: Raw blots of (a-e) pGSK3 and (f-j) GSK3β expression in the kidney organoids over time.*

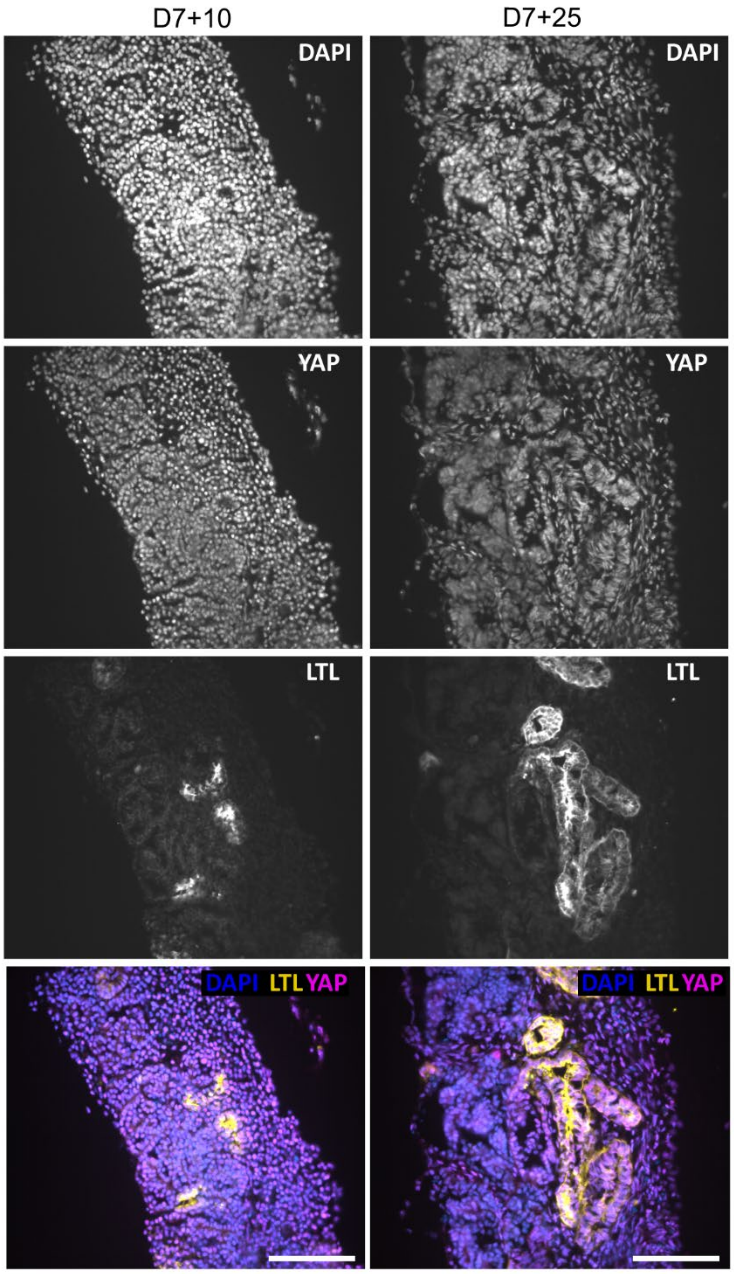


*Figure S5: YAP expression in kidney organoids at D7 + 5 and D7 + 25. Proximal tubules are stained with lotus tetragonolobus lectin (LTL). There are no obvious differences in YAP expression in podocytes (P) between D7 + 5 and D7 + 25. Scale bars: 100 µm.*

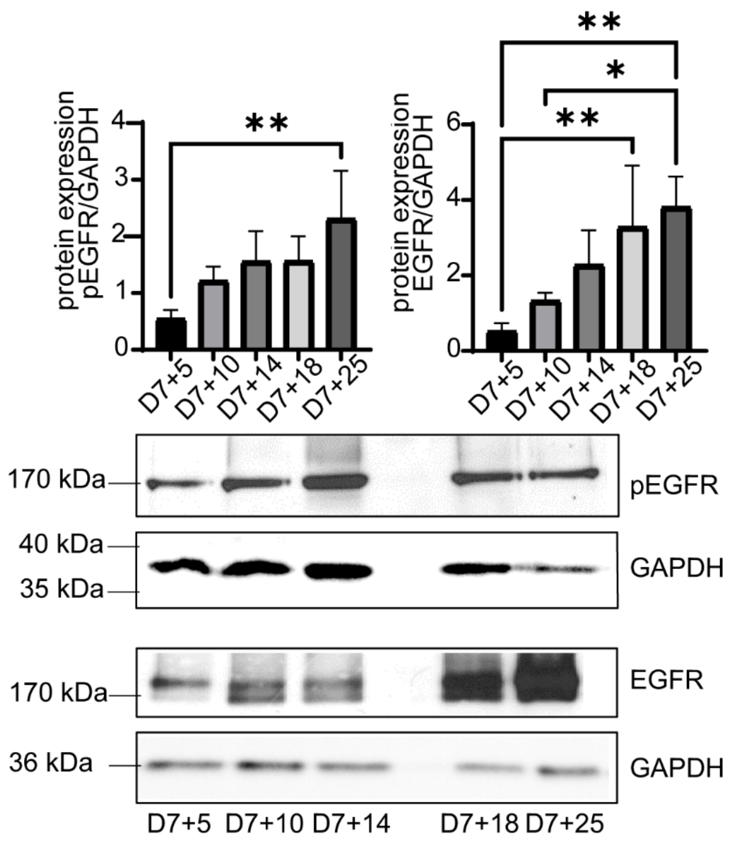


*Figure S6: Protein expression of epidermal growth factor receptor (EGFR) and phosphorylated epidermal growth factor receptor (pEGFR) over time in kidney organoids. Both pEGFR (*$p < 0.0042$*) and EGFR (*$p < 0.0008$*) were significantly upregulated over time. *:* $p = 0.013$, **: $p < 0.005$.

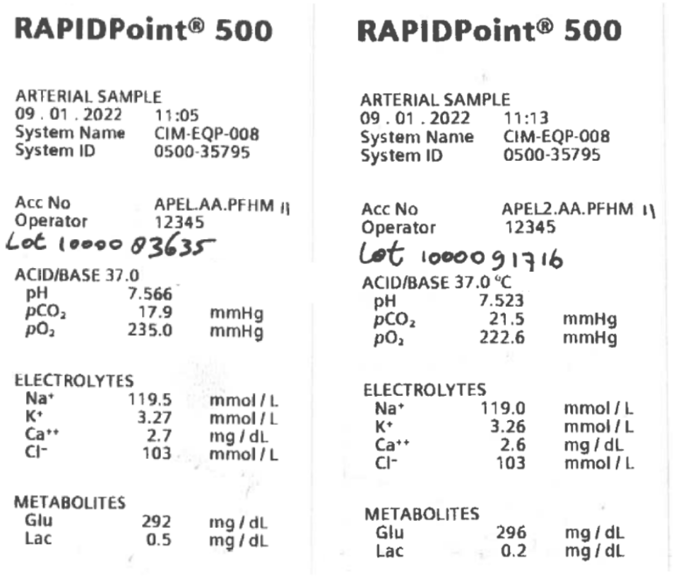

**RAPIDPoint® 500**

ARTERIAL SAMPLE
09 . 01 . 2022 11:05
System Name CIM-EQP-008
System ID 0500-35795

Acc No APEL.AA.PFHM II
Operator 12345
Lot 1000083635

ACID/BASE 37.0

| | | |
|---|---|---|
| pH | 7.566 | |
| $pCO_2$ | 17.9 | mmHg |
| $pO_2$ | 235.0 | mmHg |

ELECTROLYTES

| | | |
|---|---|---|
| $Na^+$ | 119.5 | mmol / L |
| $K^+$ | 3.27 | mmol / L |
| $Ca^{++}$ | 2.7 | mg / dL |
| $Cl^-$ | 103 | mmol / L |

METABOLITES

| | | |
|---|---|---|
| Glu | 292 | mg / dL |
| Lac | 0.5 | mg / dL |

**RAPIDPoint® 500**

ARTERIAL SAMPLE
09 . 01 . 2022 11:13
System Name CIM-EQP-008
System ID 0500-35795

Acc No APEL2.AA.PFHM 1\
Operator 12345
Lot 1000091716

ACID/BASE 37.0 °C

| | | |
|---|---|---|
| pH | 7.523 | |
| $pCO_2$ | 21.5 | mmHg |
| $pO_2$ | 222.6 | mmHg |

ELECTROLYTES

| | | |
|---|---|---|
| $Na^+$ | 119.0 | mmol / L |
| $K^+$ | 3.26 | mmol / L |
| $Ca^{++}$ | 2.6 | mg / dL |
| $Cl^-$ | 103 | mmol / L |

METABOLITES

| | | |
|---|---|---|
| Glu | 296 | mg / dL |
| Lac | 0.2 | mg / dL |

*Figure S7: Measurement of two LOTs of supplemented APEL2 organoid culture medium on a RapidPoint500 analyzer (Siemens). The glucose concentration is 294 mg/dL on average.*
