## Supplementary Methods and Tables for "Ultrastructural comparison of human kidney organoids and human fetal kidneys reveals features of hyperglycemic culture"

### Human embryonic/fetal kidneys

The human embryonic and fetal material was provided by the Joint MRC/Wellcome Trust (grant# MR/R006237/1) Human Developmental Biology Resource (www.hdbr.org). The kidneys were derived from legal abortuses and the HDBR ensured informed consent of the mother. Kidney diseases were excluded by screening the medical files. The kidneys of each embryo/fetus (Supplementary Table 1) were cut in half (longitudinally) to fix one half for histology and the other half for TEM. When the kidneys were too small, whole kidneys were fixed for either analysis. The kidneys were confirmed normal by a pathologist (T.N.) and technician (R.B.) of the University Medical Center Utrecht (the Netherlands) based on histological stainings before analysis using TEM. Details on the tissues can be found in Supplementary Table 1. A minimum of two cubes were viewed per patient.

*Supplementary Table 1: Patient information. Post-conceptional week (PCW).*

| **Embryo ID** | **Gestational age** | **Race** |
|---|---|---|
| 15333 | PCW 9 | unknown |
| 15337 | PCW 10 | white, British |
| 15342 | PCW 11 | white, British |
| 15315 | PCW 12 | unknown |
| 15317 | PCW 12 | white, British |
| 15332 | PCW 12 | white, British |
| 15335 | PCW 12 | white, British |

Induced pluripotent stem cell culture, differentiation and organoid culture

Induced pluripotent stem cells (iPSCs) were differentiated towards kidney tissue and the formed organoids were cultured as described in our previous work.(13) Briefly, the iPSC line LUMC0072iCTRL01 (male fibroblasts reprogrammed using RNA Simplicon reprogramming kit, Millipore) was passaged biweekly and maintained in Essential 8 Medium (Thermo Fisher Scientific). To start differentiation, 80,000 cells per well were seeded into a six well plate with APEL2 medium supplemented with 1% protein free hybridoma medium (PFHMII, Thermo Fisher Scientific), 1% antibiotic-antimycotic (Thermo Fisher Scientific), 8 μM GSK-3 inhibitor CHIR99021 (R&D Systems), 200 ng/ml fibroblast growth factor 9 (FGF9; R&D Systems), and 1 μg/ml heparin (Sigma-Aldrich) over the course of 7 days (D7). The cells were then aggregated and further maintained on polyester membrane inserts — either 0.4 μm pore inserts (Corning) or 1 μm pore, translucent inserts (CellQart) — in tissue culture plates. For an additional 5 days (stated as D7 + 5), the organoids were cultured with the aforementioned growth factors and small molecules for further differentiation. From D7 + 5 onwards, the organoids were cultured in basic organoid culture medium (APEL2 + 1% PFHMII + 1% antibiotic-antimycotic). The medium was refreshed every 2 days. The organoids were assessed at the following time points: D7 + 5, D7 + 10, D7 + 14, D7 + 18, and D7 + 25. In total, five independent cultures were performed (N=5).

Embedding, processing and TEM imaging

The organoids were washed once with 0.1 M cacodylate (pH 7.4), 1% sucrose and 3× with 0.1 M cacodylate (pH 7.4), followed by incubation in 1% osmium tetroxide and 1.5% $K_4Fe(CN)_6$ in 0.1 M sodium cacodylate (pH 7.4) for 1 h at 4 °C. Next, the samples were washed with MilliQ and dehydrated in a graded ethanol series (70%, 90%, 100%) at room temperature (RT). Epon infiltration followed by embedding in epon were completed with polymerization (48 h at 60 °C). The regions of interest (ROIs) were determined in toluidine blue–stained, semi-thin (1 µm) sections. Accordingly, ultrathin sections (60 nm) were cut on a Leica UC7 ultramicrotome with a diamond knife (Diatome). The sections were transferred onto 50 mesh copper grids covered with a Formvar and carbon film and subsequently stained with 2% uranyl acetate in 50% ethanol and lead citrate.

Imaging was performed with a Tecnai T12 electron microscope equipped with an Eagle 4k × 4k CCD camera (Thermo Fisher Scientific). The ROIs chosen in toluidine blue sections were targeted in the TEM by making large scans with the EPU software (1.6.0.1340REL, FEI). Subsequently, the ROI was imaged in tiles at 2900× magnification using the Mesh software (version 6). Omero and PathViewer were used to stitch, upload and annotate the data. For three cultures (N=3), one organoid per timepoint (n=1) was embedded and analyzed with TEM.

Immunofluorescence

The organoids were fixed at D7 + 5, D7 + 10, D7 + 14, D7 + 18, and D7 + 25 in 2% (v/v) paraformaldehyde (20 min, 4 °C). Cryopreservation and cutting, as well as immunofluorescence staining were performed as described previously.(13) Briefly, the cryosections were blocked with phosphate-buffered saline (PBS) containing 0.2% (v/v) Tween, 10% (w/v) bovine serum albumin (BSA) and 0.1 M glycine (20 min, RT) and incubated with primary antibodies (Supplementary Table 2) diluted in the dilution buffer of 0.2% (v/v) Tween, 1% (w/v) BSA and 0.1 M glycine in PBS (overnight, 4 °C). After washing with 0.2% (v/v) Tween in PBS, the slides were incubated with the appropriate secondary antibodies (Supplementary Table 2) diluted in the dilution buffer (1 h, RT). Next, the slides were washed with 0.2% (v/v) Tween in PBS and mounted with Prolong Gold (Thermo Fisher Scientific). The slides were cured for 2 days before imaging, which was performed on the automated Nikon Eclipse Ti-E microscope equipped with a 20× air and a 40× oil objective. All images were processed using Fiji(15), without background removal or similar modifications. For five cultures (N=5), one organoid per timepoint (n=1) was assessed.

*Supplementary Table 2: Primary and secondary antibodies and stains used for this research.*

| Target | Host | Clonality | Dilution Western blot | Dilution immuno-fluorescence | Supplier | Cat. no. |
|---|---|---|---|---|---|---|
| **Primary antibodies and stains** | | | | | | |
| B catenin | Rabbit | Recombinant (D10A8H2) | 1/300 | 1/300 | Cell Signaling Technologies | #8480 |
| GSK3β | Rabbit | Recombinant (D5C5Z) | 1/1000 | 1/400 | Cell Signaling Technologies | #12456 |
| Lotus tetragonolobus lectin (LTL), biotinylated | N/A | N/A | N/A | 1/300 | Vectorlabs | B-1325-2 |
| Nephrin | Sheep | Polyclonal | N/A | 1/300 | R&D Systems | AF4269 |
| Phospho-GSK3β | Rabbit | Recombinant (5G4) | 1/1000 | 1/200 | Cell Signaling Technologies | #9323 |
| WT1 | Mouse | Monoclonal | N/A | 1/300 | Santa Cruz Biotechnology | sc-7385 |
| YAP | Mouse | Monoclonal | N/A | 1/200 | Santa Cruz Biotechnology | sc-376830 |
| EGFR | Mouse | Monoclonal | 1/500 | N/A | Thermo Fisher Scientific | MA5-13269 |
| Phospho-EGFR | Rabbit | Monoclonal | 1/1000 | N/A | Thermo Fisher Scientific | 44-788G |
| **Secondary antibodies and stains** | | | | | | |
| Alexa anti-mouse 647 | Goat | Polyclonal | N/A | 1/400 | Thermo Fisher Scientific | A-21235 |
| Anti-mouse-HRP | Goat | Polyclonal | 1/3000 | N/A | Bio-Rad Laboratories | 1706516 |
| Alexa anti-rabbit 488 | Donkey | Polyclonal | N/A | 1/400 | Thermo Fisher Scientific | A-21206 |
| Alexa anti-rabbit 568 | Donkey | Polyclonal | N/A | 1/400 | Thermo Fisher Scientific | A10042 |
| Alexa anti-sheep 488 | Donkey | Polyclonal | N/A | 1/400 | Thermo Fisher Scientific | A-11015 |
| Alexa anti-sheep 568 | Donkey | Polyclonal | N/A | 1/400 | Thermo Fisher Scientific | A21099 |
| Streptavidin 568 | N/A | N/A | N/A | 1/400 | Thermo Fisher Scientific | S11226 |
